# Proteogenomic Characterization of Triple-Negative Apocrine Carcinoma Reveals Molecular Features of Progression and Chemotherapy Response

**DOI:** 10.1101/2024.07.30.605782

**Authors:** Yiying Zhu, Mengping Long, Wenhao Shi, Tianlong He, Fangzhou Xie, Annan Qian, Yuqiao Liu, Taobo Hu, Shaojun Tang

## Abstract

Triple-negative apocrine carcinoma (TNAC) is an exceptionally rare, chemotherapy-insensitive subtype of triple-negative breast cancer (TNBC) characterized by androgen receptor (AR) positivity and low proliferative activity. Despite poor chemotherapy response, TNAC exhibits favorable long-term survival, suggesting distinct resistance mechanisms. Here, we integrate mass spectrometry-based proteomics and whole-exome sequencing (WES) to characterize TNAC’s molecular features. We identify progressive overexpression of PI3K/AKT and AR signaling proteins, along with ECM remodeling and upregulation of GTPase-related proteins, highlighting TNAC’s invasive potential. WES reveals somatic mutations in ECM-related genes (COL18A1) and immune-associated genes (C3, NRDC), implicating the tumor microenvironment. Unlike TNBC, post-chemotherapy TNAC proteomics shows mixed ribosomal regulation and heightened inflammation, rather than reliance on proliferation, suggesting a need for targeted metabolic and immune therapies. Our findings support PI3K/AKT inhibitors and AR antagonists, potentially alongside GTPase inhibitors, metabolic disruptors, and immune therapy, as alternative strategies to standard chemotherapy in TNAC.

## Introduction

Triple-negative apocrine carcinoma (TNAC) is a rare and chemo-insensitive subtype of breast cancer. While apocrine carcinoma (AC) accounts for 0.4–4% of all breast cancers, TNAC, representing approximately 1% of triple-negative breast cancers (TNBCs), is estimated to have around one thousand new cases annually in China, making it one of the least studied subtypes^1–5^. TNBC is defined by the absence of estrogen receptor (ER), progesterone receptor (PR), and Human Epidermal Growth Factor Receptor 2/ Erb-B2 Receptor Tyrosine Kinase 2 (ERBB2, also known as HER2), which limits targeted treatment options and often leads to aggressive clinical behavior^5,6^. Despite its rarity, TNAC differs from non-apocrine triple-negative breast cancer (NA-TNBC) by exhibiting low Ki67 expression and androgen receptor (AR) positivity, which may contribute to a lower proliferative index and distinct therapeutic vulnerabilities^7–9^.

AC is characterized by apocrine morphology, including large tumor cells, eosinophilic cytoplasmic granules, and prominent nucleoli^10,11^. It can present as ductal carcinoma in situ (DCIS) or invasive carcinoma. Originating from epithelial cells within the terminal ductal lobular units, DCIS remains confined within the ducts, while invasive breast cancer penetrates the basement membrane and stroma, requiring systemic therapies to prevent or manage metastasis. According to the World Health Organization, invasive apocrine carcinoma is defined by tumors in which over 90% of cells exhibit apocrine morphology^12^. Understanding the molecular mechanisms driving this progression is crucial for developing effective treatment strategies.

TNAC exhibits poor response to chemotherapy yet better long-term survival, sparking interest in personalized treatment approaches^4,7,9,13–16^. While multidrug chemotherapy remains a standard treatment for malignant tumors, neoadjuvant chemotherapy—administered before surgery to shrink tumors—is increasingly utilized^6,17,18^. However, its side effects and variable efficacy have led to a growing focus on identifying patients who may not benefit from chemotherapy^19,20^. In ER+/PR+ and HER2+ breast cancers, treatment decisions are carefully guided by chemo-exemption strategies^21–24^. In TNBC, immunotherapy (e.g., pembrolizumab) and AR-targeted therapies are emerging alternatives^25–28^. Increasing evidence suggests that TNAC patients may not benefit from chemotherapy, prompting discussions on chemo-exemption to optimize treatment and reduce unnecessary toxicity^9,16^.

Our team conducted a retrospective clinical and statistical analysis of 41 TNAC cases and the SEER database, revealing that while TNAC exhibits a poor response to neoadjuvant therapy, it has a better short-term prognosis than other triple-negative breast cancers, likely due to its low proliferative nature^9^. However, the molecular mechanisms underlying these clinical observations remain unexplored. Previous studies have primarily focused on genomic profiling to classify TNAC subtypes and identify potential drug targets^29,30^, yet a comprehensive proteogenomic approach is still lacking. To address this gap, this study integrates mass spectrometry (MS)-based proteomics and whole-exome sequencing (WES) to systematically characterize dysregulated proteins, altered pathways, and somatic mutations in TNAC. We constructed a proteome landscape to identify key molecular features of TNAC progression and examined somatic mutations to assess their impact on protein expression dysregulation. Additionally, we evaluated the chemotherapy response of TNAC tumors, providing insights into the molecular basis of chemotherapy resistance. Overall, this study offers a comprehensive proteogenomic perspective on TNAC, contributing to the development of precision medicine strategies tailored to this rare subtype.

## Results

### Clinical and pathological characteristics of samples

In order to explore TNAC as a unique tumor type, we collected formalin-fixed paraffin-embedded (FFPE) samples from 31 patients diagnosed with invasive TNAC (Supplementary Table 1). The general workflow is pictured in Figure 1a. The median age of patients at diagnosis was 57 years, ranging from 38 to 79. 41.9% (13/31) of patients were at TNM stage I, 45.2% (14/31) were at stage II, and 12.9% (4/31) were at stage III. The morphology of samples was evaluated and confirmed by two pathologists, and protein markers were confirmed by immunohistochemical staining (IHC). Example IHC and Hematoxylin and Eosin (H&E) images are shown in Figure 1b/c. All patients tested negative for the hormone receptors ER and PR, and were non-amplified for HER2. They had positive AR and low Ki67, along with an increasing Epidermal growth factor receptor (EGFR) from normal to invasive tissues. The average duration of storage for FFPE samples was 1869 days, ranging from 229 to 3185 days. Ten patient samples still contain DCIS tumor tissues. Seven patients received neoadjuvant chemotherapy before surgery, and their needle biopsy FFPEs were also collected. Among 7 neoadjuvant chemotherapy patients, 3 were at TNM stage II, 2 were at stage III, and 1 was at stage I, with a median age of 54 (from 46 to 63). Pathological responses to neoadjuvant chemotherapy were assessed using the Miller–Payne (MP) grading system on a five-point scale. Four patients had an MP grade 2 response, while 3 had a grade 3 response. In conclusion, a total of 78 samples were collected in the study, including 30 adjacent tissues (named as normal tissues), 10 DCIS tissues, and 38 invasive tissues (from 31 patients, including 7 pre-chemotherapy biopsies). All FFPE tissues were analyzed by MS, while invasive and normal tissues from three patients were analyzed by WES.

**Figure 1:**
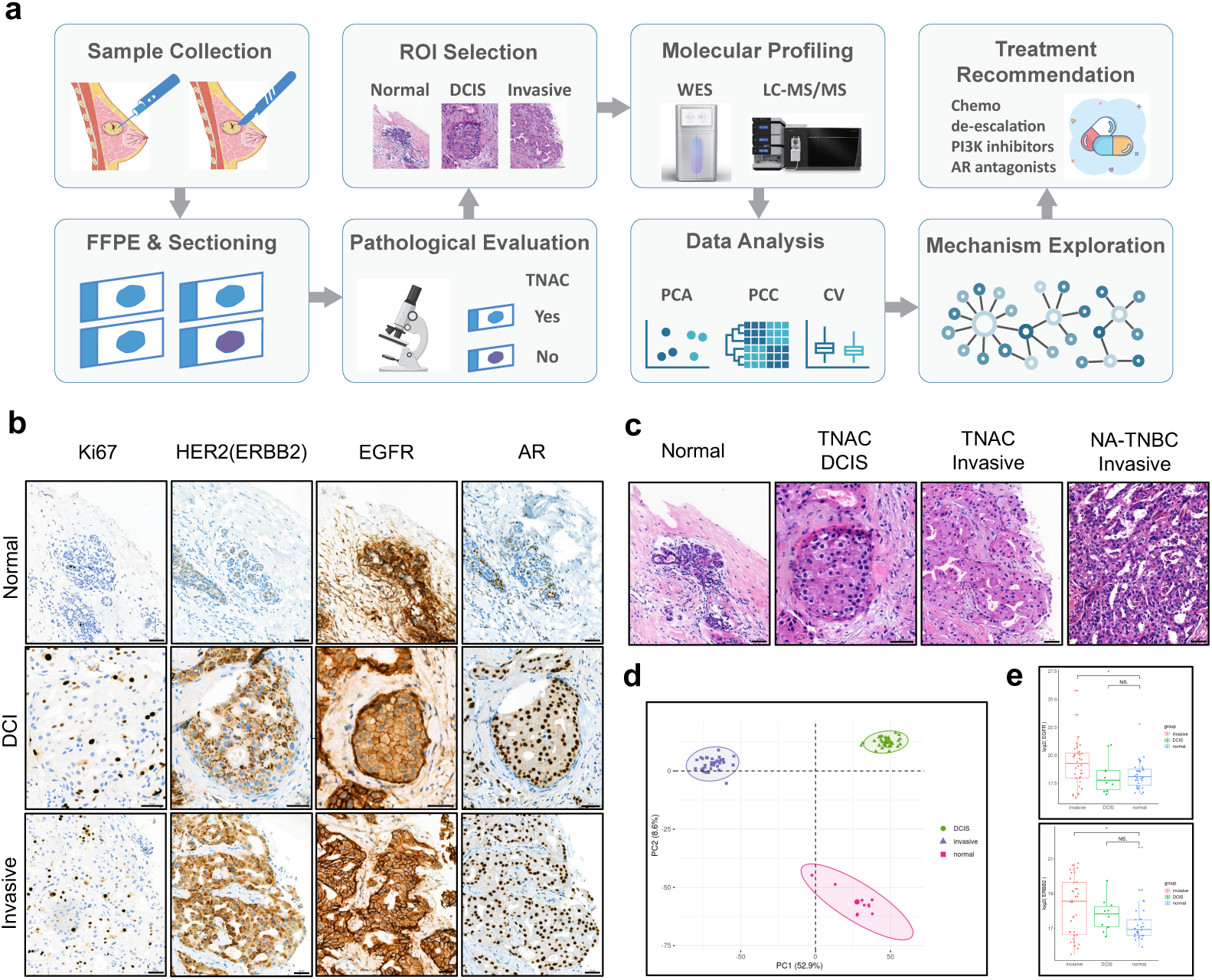
Overview of the experimental design and representative H&E and IHC images of clinical samples. (a) Schematic workflow of the study. Breast cancer patient samples were collected from needle biopsies before neoadjuvant chemotherapy and tumor resections during surgery. FFPE sections were reviewed by pathologists, and TNAC classification was confirmed. Normal, DCIS, and invasive tumor tissues were dissected for proteomic analysis using LC-MS/MS, with selected samples undergoing whole-exome sequencing (WES). The data were then processed for bioinformatic analysis. (b) Representative H&E-stained images of normal breast tissue, TNAC DCIS, TNAC invasive carcinoma, and NA-TNBC invasive carcinoma. TNAC tumors exhibit characteristic apocrine morphology, including abundant eosinophilic cytoplasm, distinct nucleoli, and well-defined cell boundaries. Scale bar = 50 μm. (c) Immunohistochemical (IHC) staining of Ki67, HER2 (ERBB2), EGFR, and AR in normal, DCIS, and invasive TNAC tissues. Scale bar = 50 μm. (d) Principal component analysis (PCA) (left) and representative protein expression data from LC-MS/MS (right). PCA clustering shows a clear separation of normal, DCIS, and invasive TNAC tissues, indicating distinct molecular profiles. (e) Boxplots displaying MS-based expression levels of EGFR (top) and HER2/ERBB2 (bottom) across normal, DCIS, and invasive TNAC tissues (\**p* < 0.05).

Current MS-based proteomics analysis quantified 5952 protein groups. TNAC typically expresses protein AR and EGFR while lacking ER, PR, and Ki67^7,9,11,13,14^. Our MS data on protein expression aligns with the expectation and the observation from IHC (Fig. 1b/d). Drawn by this TNAC proteomics data analysis, we found that ER, PR, and Ki67 were not detectable in tumor tissues, while EGFR was strongly expressed in invasive tissues, and AR was also present in invasive tissues. Notably, protein HER2 was low expressed, as observed in IHC images and MS data (Fig. 1b/d). Nevertheless, it is possible that MS fails to detect relatively low-abundant proteins due to its instrumentation^31^. For example, GCDFP-15 was not detected in our proteomics data but was detected in IHC. Overall, the conclusion drawn from proteomics analyses is consistent with the IHC study and previous reports.

### Proteomic and Functional Analysis of TNAC Progression

Understanding the molecular characteristics can help us grasp the initiation and progression of the tumor. Invasive and DCIS tissues, along with adjacent normal tissues, were carefully removed from selected TNAC FFPE samples and underwent MS-based proteomics analysis. The representative IHC and H&E images are shown in Figure 1b/c. In total, 5952 protein groups were quantified in this dataset, with 4065 proteins from normal tissues, 3844 proteins from DCIS tissues, and 5277 proteins from invasive tissues (Supplementary Table 2). Principal component analysis (PCA) (Fig. 1d) demonstrated distinct clustering between normal, DCIS, and invasive TNAC tissues, reflecting significant molecular divergence. Differential protein expression and heatmap analysis identified 2,309 proteins with significant alterations, highlighting a progressive molecular shift from DCIS to invasive TNAC (Fig. 2a; Supplementary Table 3).

**Figure 2:**
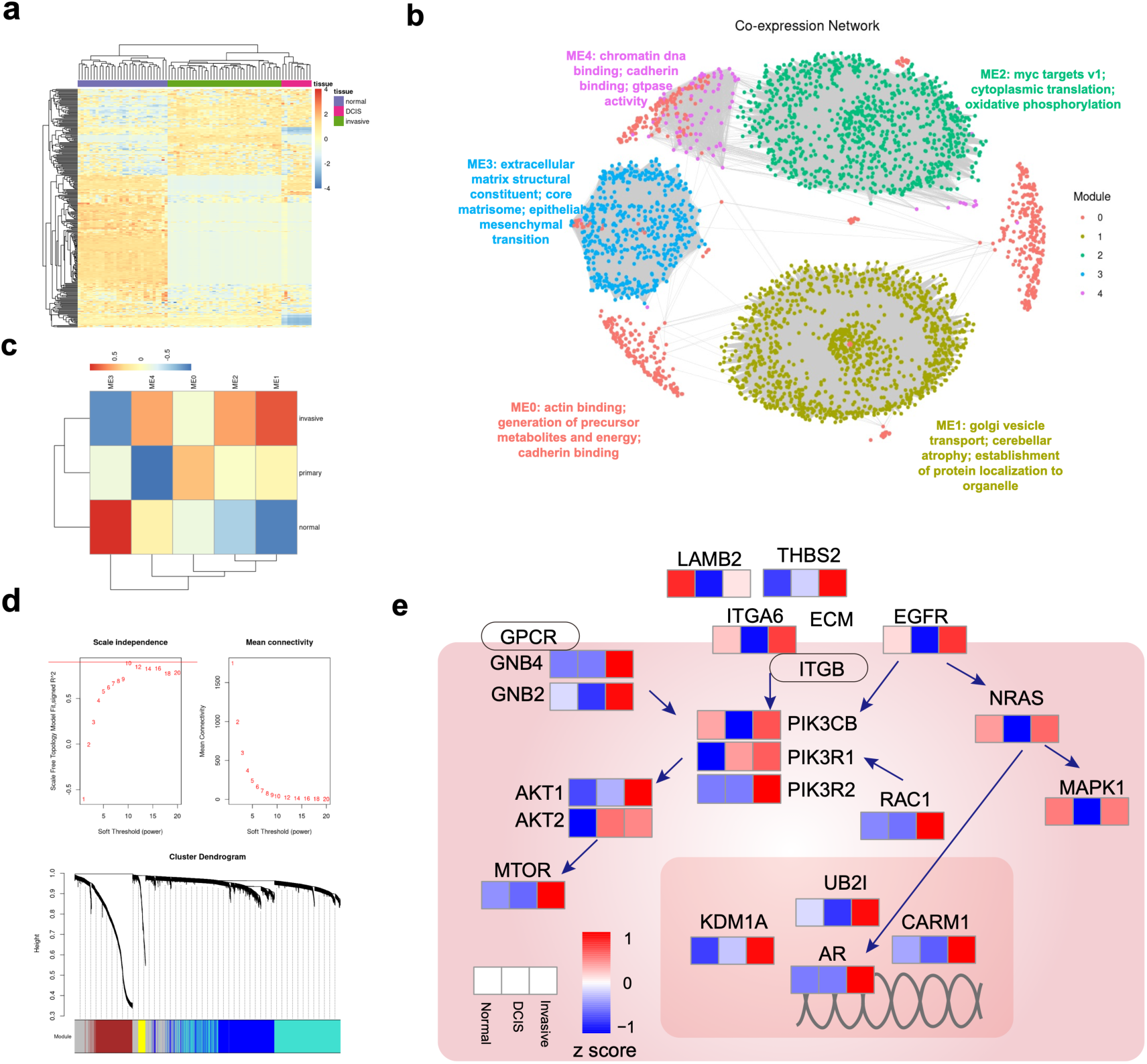
Proteomic and network analysis of TNAC tumor progression. (a) Heatmap showing variations in protein expression across normal, DCIS, and invasive TNAC tissues. Each cell represents the standardized score of protein abundance in an individual sample, with colors indicating relative expression levels. (b) Weighted gene co-expression network analysis (WGCNA) identified five major protein co-expression modules (ME0–ME4) in TNAC samples. Each node represents a protein, with colors corresponding to functionally distinct modules. ME2 is associated with MYC targets and oxidative phosphorylation, ME3 with extracellular matrix remodeling and epithelial-mesenchymal transition (EMT), and ME4 with chromatin and cadherin binding. (c) Bilateral clustering of the five modules in normal, DCIS, and invasive TNAC tissues. (d) Construction of the WGCNA co-expression network. (Top) Selection of the soft threshold power for network scale-free topology. (Bottom) Dendrogram visualization of co-expressed protein modules. (e) Signaling pathways illustrating the upregulation of proteins involved in PI3K/AKT, androgen receptor (AR) signaling in invasive TNAC. The figure highlights the increased expression of key regulators, including PI3K, AKT, mTOR, AR, and RAC.

Weighted correlation network analysis (WGCNA) revealed key co-expression modules associated with TNAC progression (Fig. 2b). A total of 3277 proteins were grouped into five functional modules (ME0-ME4), each reflecting distinct biological processes. ME0, enriched in actin binding, suggests cytoskeletal organization, while ME1, associated with Golgi vesicle transport and protein localization, indicates altered intracellular trafficking. ME2, linked to MYC targets and oxidative phosphorylation, highlights metabolic reprogramming in TNAC. ME3, featuring extracellular matrix (ECM) remodeling and epithelial-mesenchymal transition, underscores the structural changes supporting tumor invasion. ME4, enriched in chromatin DNA-binding proteins, suggests transcription and epigenetic regulation. Bilateral clustering revealed that ME3 was most prominent in normal tissues, ME0 in DCIS, and ME1 in invasive tumors, illustrating progressive molecular shifts from tumor initiation to invasion (Fig. 2c).

Differential expression analysis identified significant changes in key biological pathways associated with TNAC progression (Fig. 3; Supplementary Table 3). A total of 1,647 proteins were upregulated, and 463 were downregulated in invasive tumors compared to normal tissues. Invasive TNAC exhibited increased cadherin binding and GTPase activity, supporting cell adhesion modulation and cytoskeletal reorganization, whereas ECM and actin-binding proteins were downregulated, indicating progressive loss of structural integrity. Comparisons between DCIS and normal tissues showed cadherin binding and ligase activity enrichment while ECM-associated proteins were downregulated, resembling patterns observed in invasive TNAC. Additionally, invasive tumors displayed enrichment in GTPase activity and guanyl ribonucleotide binding proteins, whereas DCIS retained ECM-associated proteins, suggesting different tumor microenvironment adaptations at distinct disease stages.

**Figure 3:**
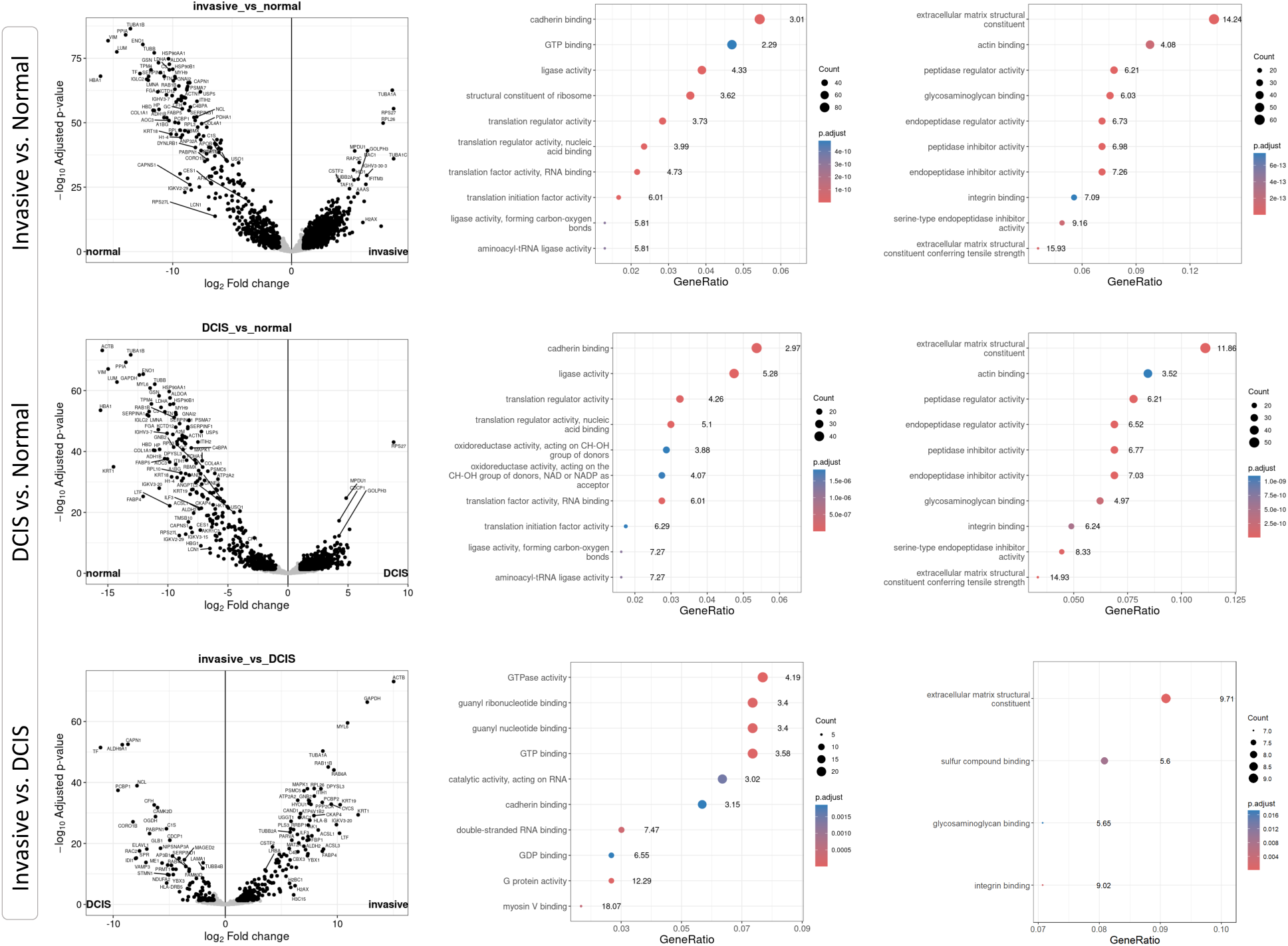
Differential expression and functional enrichment analysis of TNAC normal, DCIS, and invasive tissues. (Left) Volcano plots illustrating differentially expressed proteins across three comparisons: invasive vs. normal, DCIS vs. normal, and invasive vs. DCIS. The x-axis represents log₂ fold change, while the y-axis represents –log₁₀ p-value. Significantly altered proteins (*p* < 0.05, |log₂ fold change| > 1) are highlighted in black. (Middle) Top enriched Gene Ontology (GO) terms for upregulated proteins in each comparison, categorized by molecular functions and biological processes (*p* < 0.05, log₂ fold change > 1). The numbers beside the dots represent fold enrichment. The dot size represents the number of proteins involved, while the color gradient indicates the adjusted p-value. (Right) Top enriched GO terms for downregulated proteins in each comparison (*p* < 0.05, log₂ fold change < -1). The numbers beside the dots represent fold enrichment. The dot size corresponds to the number of proteins in each category, and the color represents statistical significance (*p*-adjusted values). The results highlight distinct molecular features of TNAC progression, including cadherin and GTPase binding in invasive tumors and extracellular matrix remodeling in DCIS.

The Gene set enrichment analysis (GSEA) analysis further highlighted key oncogenic drivers of TNAC (Supplementary Table 3). The Phosphoinositide 3-kinase (PI3K)/ Protein kinase B (PKB, also known as AKT) signaling was significantly enriched across tumor progression (*p* < 0.01), confirming its role as a primary oncogenic driver in TNAC. Similarly, AR signaling was activated in invasive TNAC vs. DCIS (*p*-value = 0.035), reinforcing its role in tumor progression and potential therapeutic relevance. The analysis also revealed metabolic and immune adaptations, including increased mitochondrial translation in invasive TNAC, dysregulated cholesterol synthesis in DCIS, and complement pathway inhibition in both tumor types, possibly contributing to immune evasion. The epigenetic regulatory processes were more active in invasive TNAC, while DCIS tumors exhibited increased activity in scavenging by class A receptors, suggesting early-stage microenvironmental adaptations. These findings suggest that targeting PI3K/AKT and AR signaling could provide therapeutic opportunities in TNAC while highlighting ECM remodeling and metabolic reprogramming as key hallmarks of tumor progression.

### PI3K/AKT and AR pathway activation in TNAC progression

The PI3K/AKT pathway has long been recognized as a key driver and therapeutic target in breast cancer^32–34^. While PIK3CA somatic mutations were less frequently detected in TNBC compared to other breast cancer subtypes^35^, certain TNBC subtypes exhibit higher PI3K mutation rates, making PI3K inhibitors a potential treatment option^33,36^. In TNAC, genomic studies have consistently identified mutations in PIK3CA, PIK3R1, PTEN, and TP53, highlighting PI3K/AKT pathway activation as a hallmark of this subtype^29,30^. Lyman et al. classified TNAC into six molecular subgroups based on gene expression, including the luminal androgen receptor (LAR) subtype, which Kim et al. found to be the most common TNAC subtype, accounting for approximately 40% of cases^29,37^. While previous genomic studies failed to detect mutations in the AR gene in TNAC^29,30^, our proteomics analysis confirms AR overexpression in invasive TNAC tissues, reinforcing its role in tumor biology.

The cross-talk between AR and PI3K/AKT signaling has been well-documented in prostate cancer^38–43^, and our findings suggest a similar interaction in TNAC. Figure 2e illustrates the overexpression of proteins involved in both the PI3K/AKT and AR pathways in invasive TNAC tissues. The upregulation of PIK3CB, PIK3R1, PIK3R2, AKT1, AKT2, and Mechanistic Target of Rapamycin (mTOR) in invasive TNAC, compared to DCIS and normal tissues, suggests enhanced proliferative and survival signaling. Additionally, increased AR expression, along with upregulation of Coactivator-associated arginine methyltransferase 1 (CARM1), Ubiquitin-Conjugating Enzyme E2 I (UBE2I), and Lysine Demethylase 1A (KDM1A), indicates active AR transcriptional regulation. Additionally, upregulated Ras-related C3 botulinum toxin substrate1 (RAC1) and Mitogen-Activated Protein Kinase 1 (MAPK1) highlight GTPase-mediated cytoskeletal reorganization, reinforcing the metastatic potential of TNAC. The concurrent activation of PI3K/AKT/mTOR and AR pathways underscores their oncogenic role, supporting therapies using PI3K inhibitors and/or AR antagonists, while GTPase inhibitors targeting RAC1 may help mitigate TNAC invasion.

### WES Analysis of TNAC Tumors

WES analysis was conducted on six paired invasive and normal tumor tissue samples. Figure 4 provides an overview of the WES sequencing results. The differential analysis yielded 32 mutated genes, with splice sites being the dominant mutation type, followed by in-frame insertions and deletions (Fig. 4a). The most common single nucleotide variation (SNV) class was C to T transitions (Fig. 4b). Through genomic sequencing, the top ten mutated genes, including NRDC, were identified (Fig. 4c). Interestingly, our analysis did not detect PI3K somatic mutations, which are frequently reported in TNAC. This absence could be attributed to individual genetic variability or sample preservation limitations in FFPE tissues.

**Figure 4.**
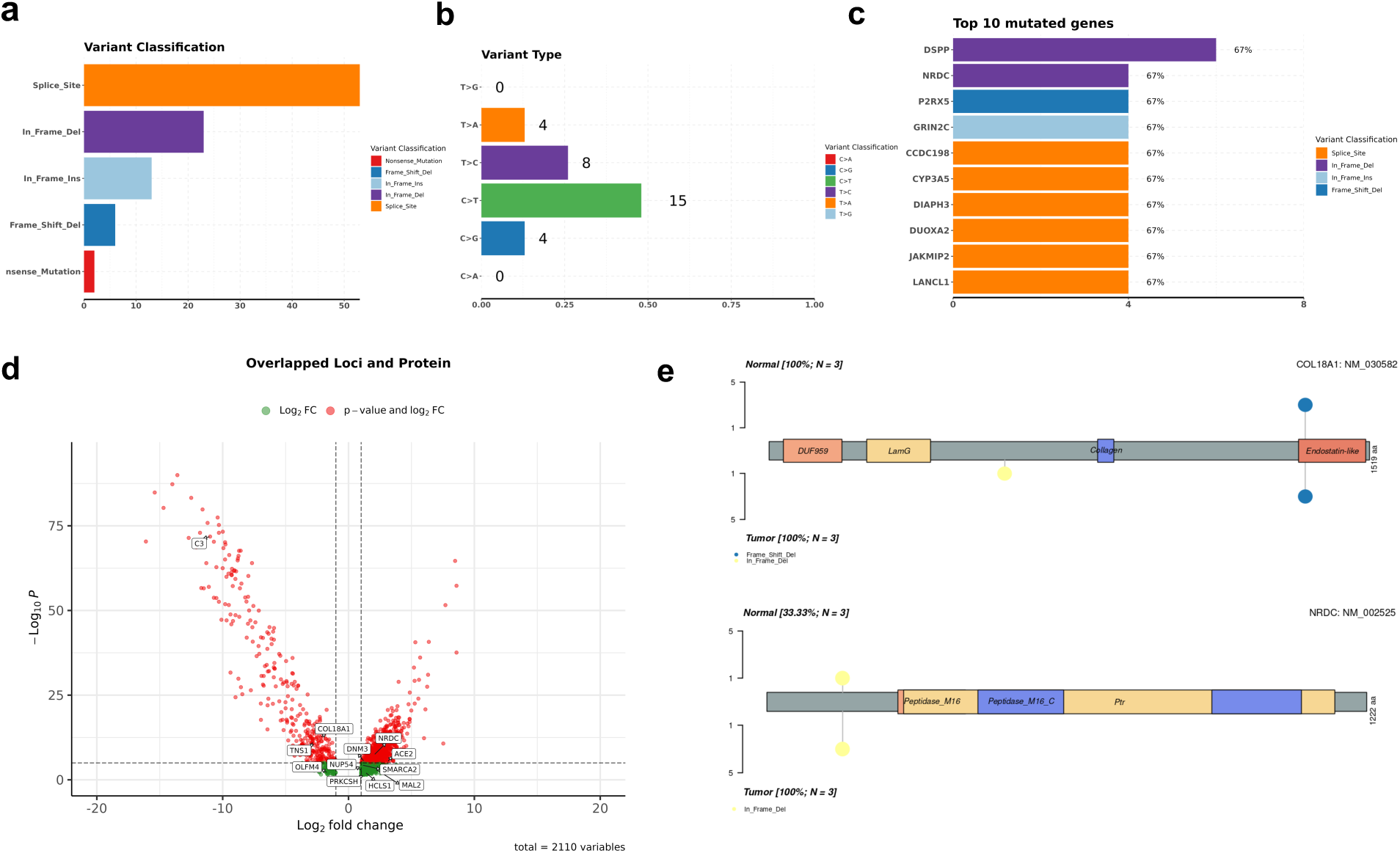
Whole-exome sequencing (WES) analysis of six paired invasive and adjacent normal TNAC samples. (a) Variant classification of identified mutations, with splice site variants being the most frequent. (b) Single nucleotide variant (SNV) type distribution, showing that the predominant mutation type is C>T. (c) Top 10 most frequently mutated genes, with bars representing the number of mutations and the percentage of TNAC patients harboring them. (d) Integration of WES and proteomics data, highlighting 12 genes that are both mutated and exhibit significant differential protein expression (*p* < 0.01, |log₂ fold change| > 1). (e) Schematic representation of mutations in COL18A1 and NRDC, illustrating the mutation sites (lollipops) detected in TNAC tumors. Colors indicate different mutation classifications, and the length of each stick represents the number of patients with the mutation.

We further compared these loci with proteomic results and identified twelve overlapping loci and their corresponding proteins: ACE2, C3, COL18A1, DNM3, HCLS1, MAL2, NRDC, NUP54, OLFM4, PRKCSH, SMARCA2, TNS1 (Fig. 4d). Among these genes, C3 exhibited the most significant changes in protein expression, suggesting that its gene mutation may have cis-effects on the protein and play crucial roles in the regulation of TNAC. Detailed analysis of the mutations in the overlapping genes revealed that C3, ACE2, MAL2, OLFM4, and DNM3 had splice site mutations; TNS1, HCLS1, and NUP54 had nonsense mutations; COL18A1 had frameshift deletions and NRDC, PRKCSH and SMACR2 have in-frame deletion. Example mutation sites are shown in Figure 4e.

Complement 3 (C3) has been reported to be a key molecule in the activation of B lymphocytes^44^. Therefore, the mutation of C3 gene and the reduction of C3 protein level may indicate that there is a disorder in B cell activation and immune suppression in this group of patients. The gene of Collagen Type XVIII Alpha 1 (COL18A1) has been reported to be significantly upregulated in human breast cancer and strongly associated with poor prognosis in high-grade breast cancer, and ablation of COL18A1 in breast cancer significantly improves the efficacy of HER2-targeted therapy^45^. COL18A1 gene is mutated in TNAC, and its protein expression is reduced in invasive tissues (Fig. 4d). Nardilysin (NRDC) was identified as a frequently mutated gene in TNAC, with an in-frame deletion detected in tumors. As a zinc metalloendopeptidase, NRDC regulates protein shedding, transcriptional control, and immune signaling, and prior studies link NRDC to tumor progression and cytokine release, suggesting its role in modulating the tumor microenvironment^46,47^. The mutation observed in TNAC may disrupt these functions, potentially contributing to immune evasion and proteomic dysregulation, warranting further investigation as a biomarker or therapeutic target.

In summary, WES results identified key mutated genes and integrated analysis with protein expression differences, providing potential mechanism clues for the occurrence and development of breast cancer from the perspectives of cellular, extracellular, and cellular immunity.

### Molecular Adaptations in Chemotherapy-Resistant TNAC

This study investigates the molecular impact of chemotherapy on intrinsically resistant cancer tissue, focusing on proteomic changes before and after treatment in patients known to exhibit poor chemotherapy response yet high overall survival rates. First, we did a PCA plot and heatmap analysis on proteomic profiles of four groups: pre-chemotherapy normal, pre-chemotherapy invasive, post-chemotherapy normal, and post-chemotherapy invasive (Fig. 5a/b). The distinct clustering patterns indicate substantial differences between normal and invasive samples, as well as pre- and post-chemotherapy states. The ellipses suggest that post-chemotherapy invasive samples cluster more closely together, whereas pre-chemotherapy normal samples exhibit a broader distribution, reflecting a higher degree of heterogeneity. Then, we focused on pre- and post-chemotherapy invasive samples. The Pearson Correlation Coefficient (PCC), PCA, and heatmap are consistent (Fig. 5c/d/e), showing that post-chemotherapy invasive samples form a more cohesive cluster, whereas pre-chemotherapy samples exhibit more variability. This suggests that chemotherapy alters the proteomic landscape of invasive tumors, potentially driving a more uniform molecular state. Differential analysis shows that 306 proteins were downregulated, 334 proteins were upregulated, while 3238 proteins were unchanged (Fig. 5f; Supplementary Table 4).

**Figure 5.**
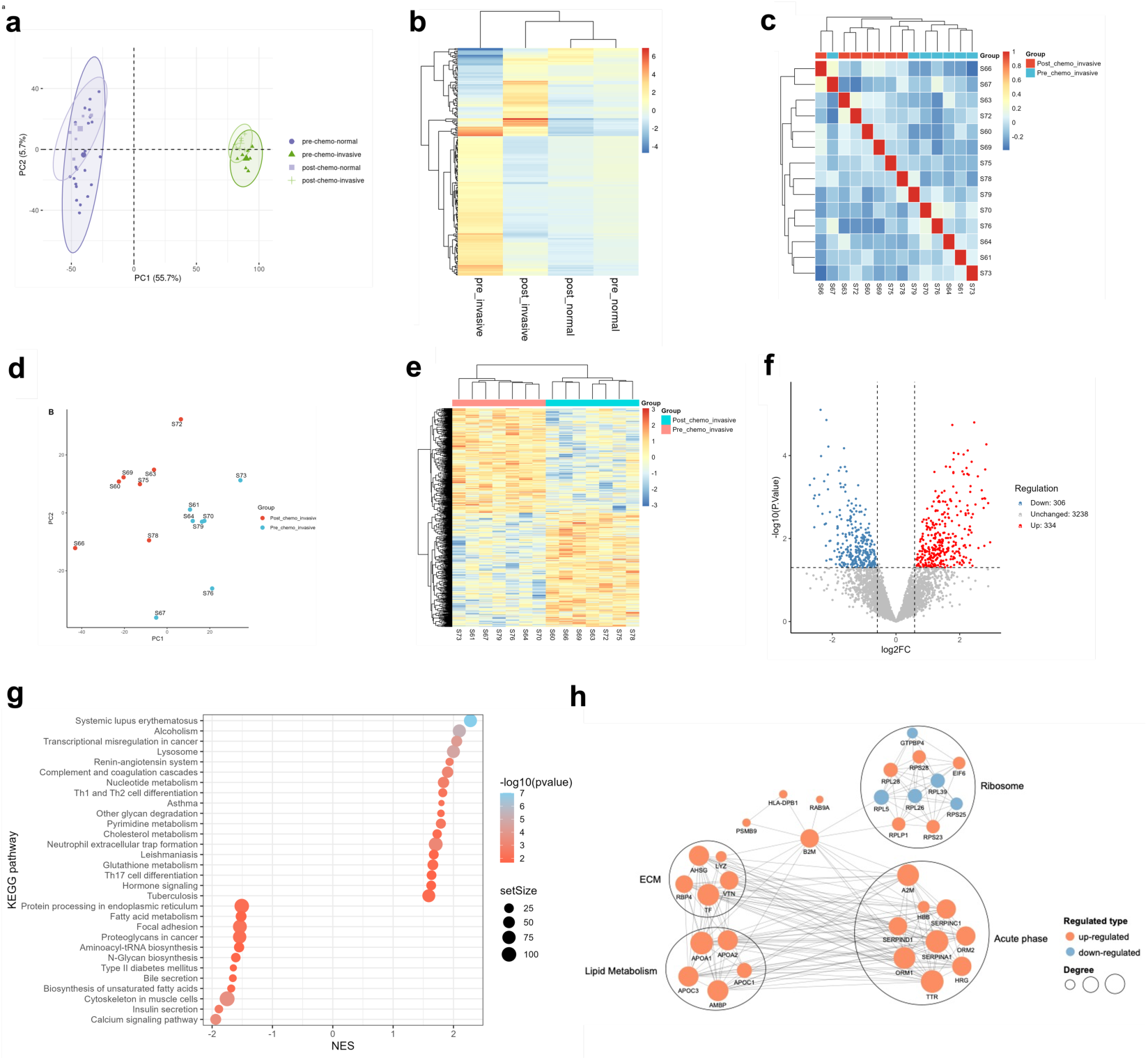
Proteomic profiling and functional analysis of pre- and post-chemotherapy TNAC samples. (a) Principal component analysis (PCA) plot showing distinct clustering of pre- and post-chemotherapy samples, with separation between normal and invasive tissues. (b) Heatmap illustrating global protein expression changes between pre- and post-chemotherapy groups. Each column represents a sample, and the color scale indicates relative protein abundance. (c) Pearson Correlation Coefficient (PCC) heatmap demonstrating hierarchical clustering of pre- and post-chemotherapy invasive samples. (d) PCA of invasive samples pre- and post-chemotherapy. (e) Heatmap of differentially expressed proteins between pre- and post-chemotherapy invasive samples, illustrating significant changes in protein expression patterns. (f) Volcano plot displaying significantly upregulated (red) and downregulated (blue) proteins in post-chemotherapy invasive samples compared to pre-chemotherapy samples (*p* < 0.05, |log₂ fold change| > 1). (g) KEGG pathway enrichment analysis of differentially expressed proteins, showing significantly upregulated and downregulated pathways in post-chemotherapy samples. (h) Protein-protein interaction (PPI) network of key functional categories enriched in post-chemotherapy samples. Nodes represent proteins, with red indicating upregulation and blue indicating downregulation.

The KEGG pathway enrichment analysis revealed significant differences in biological processes between post- and pre-chemotherapy TNAC samples, particularly in immune response, metabolism, and cellular structure regulation (Fig. 5g; Supplementary Table 4). Notably, pathways related to immune activation, such as systemic lupus erythematosus, complement and coagulation cascades, and neutrophil extracellular trap formation, were significantly upregulated after chemotherapy, suggesting an increased inflammatory response and immune system engagement. In contrast, pathways associated with metabolism, such as fatty acid metabolism, bile secretion, and calcium signaling, were downregulated post-chemotherapy, indicating metabolic shifts that may reflect altered tumor cell adaptability or stress responses. Moreover, the suppression of focal adhesion and cytoskeletal organization pathways suggests potential chemotherapy-induced effects on tumor cell migration and invasion.

The PPI analysis further supports these findings by showing distinct alterations in functional protein clusters post-chemotherapy (Fig. 5h). Ribosomal proteins (RPS23, RPS25, RPS28, RPL5, RPL26, RPL28, RPLP1, RPL39, EIF6, GTPBP4) exhibited mixed regulation, indicating selective changes in protein synthesis machinery, which may reflect cellular stress responses or adaptive resistance mechanisms. In contrast, lipid metabolism proteins (APOA1, APOA2, APOC1, APOC3, AMBP) and acute-phase response proteins (A2M, ORM1, ORM2, SERPINA1, SERPINC1, SERPIND1, HRG, HBB, TTR) were consistently upregulated post-chemotherapy. This suggests a potential increase in inflammatory and stress response proteins, reinforcing the KEGG findings on immune activation. Similarly, the upregulation of ECM proteins (VTN, AHSG, RBP4, TF, LYZ) aligns with the observed enrichment of complement and coagulation cascades, indicating remodeling of the tumor microenvironment in response to treatment.

Together, these analyses suggest that post-chemotherapy TNAC tumors undergo significant immune activation, metabolic reprogramming, and ECM remodeling. While chemotherapy may induce stress responses that affect ribosomal activity and protein synthesis, it also enhances inflammatory and acute-phase responses, which could either contribute to tumor suppression or therapy resistance. Understanding these molecular adaptations is crucial for identifying potential vulnerabilities and developing targeted therapeutic strategies for TNAC. Our findings on post/pre-chemotherapy TNAC proteomics profiles align with Anurag et al. ’s study of chemotherapy resistance and response in TNBC, highlighting metabolism reprogramming as a key resistance mechanism^48^. However, unlike TNBC, which relies on DNA damage repair and enhanced translation, TNAC exhibits mixed ribosomal regulation and a stronger inflammatory response, suggesting a distinct adaptation strategy. These distinct molecular features suggest that TNAC may require a different therapeutic approach, emphasizing metabolic and immune-targeting strategies rather than conventional chemotherapy-based regimens.

## Discussion

This study provides the first comprehensive proteogenomic characterization of TNAC, identifying key molecular features of tumor progression and chemotherapy resistance. Proteomic analysis of normal tissues, DCIS, and invasive TNAC revealed progressive molecular changes, especially the increased expression of proteins associated with the PI3K/AKT and AR signaling pathways in invasive tumors. These pathways, known to drive oncogenic proliferation and survival, suggest a mechanistic shift toward growth factor-mediated and AR-driven oncogenic signaling in invasive TNAC. GO analysis revealed a strong ECM remodeling signature and increased expression of GTPase-related proteins during tumor initiation and progression. WGCNA further identified distinct protein modules linked to tumor metabolism and signaling adaptation. MYC-driven oxidative phosphorylation (ME2) and epithelial-mesenchymal transition-related ECM remodeling (ME3) were enriched in invasive TNAC, reinforcing the metabolic plasticity and structural modifications driving tumor progression. WES uncovered somatic mutations affecting ECM-related genes, including COL18A1, as well as immune regulatory mutations in C3, suggesting a role for immune evasion in TNAC persistence.

Proteomic analysis of TNAC tissues before and after neoadjuvant chemotherapy reveals distinct molecular adaptations in response to treatment. A notable paradox in TNAC is its poor chemotherapy response despite relatively favorable long-term survival. While anthracyclines, carboplatin, and taxanes remain the standard of care for TNBC^17^, invasive TNAC exhibits a unique metabolic reprogramming profile post-chemotherapy, as highlighted by PPI network analysis. Upregulation of key metabolic adaptation proteins (ORM1, ORM2, APOA2, AHSG) suggests a shift toward lipid metabolism and acute-phase response pathways rather than apoptosis, indicating that TNAC tumors evade chemotherapy through metabolic flexibility and stress-response activation rather than relying on proliferation. This aligns with their low Ki67 expression and limited sensitivity to cytotoxic agents. Unlike conventional TNBC, where chemotherapy often selects for highly proliferative clones, TNAC tumors instead exhibit enhanced metabolic adaptation and immune modulation following treatment, highlighting the need for alternative therapeutic strategies targeting metabolism and immune responses.

Given these findings, an important clinical question arises: Does chemotherapy truly benefit TNAC patients, or could alternative, less toxic treatments provide superior outcomes? The persistence of metabolic adaptation and immune-modulatory pathways post-chemotherapy suggests that TNAC tumors do not rely on rapid proliferation but instead survive through alternative mechanisms. This may explain why TNAC patients can achieve favorable long-term survival despite limited chemotherapy response. Rather than standard chemotherapy, these patients may benefit from targeted strategies that disrupt key survival pathways, such as metabolic inhibitors, immune-modulating agents, or lipid metabolism-targeting therapies. Furthermore, these findings underscore the need for biomarker-driven treatment stratification, as some TNAC tumors may remain clinically stable without requiring chemotherapy.

These findings support the potential de-escalation of chemotherapy in TNAC treatment by adopting therapeutic alternatives. PI3K inhibitors (e.g., Alpelisib, Copanlisib)^49–51^, in combined with AR-targeted therapies (e.g., Enzalutamide)^52,53^, may offer a more effective strategy than conventional chemotherapy. Additionally, the enrichment of lipid metabolism and oxidative phosphorylation pathways highlights potential vulnerabilities of oxidative phosphorylation inhibitors (e.g., IACS-010759)^54^. Furthermore, ECM remodeling and GTPase signaling activation in invasive TNAC suggest that RAC inhibitors (e.g., EHT1864)^55^ may help limit invasion and metastasis. Given the presence of immune-related mutations and post-chemotherapy immune modulation, further exploration of immune-targeting strategies, including immune checkpoint inhibitors (e.g., PD-L1 inhibitor) ^56,57^, may be warranted in TNAC. Future studies should explore whether replacing chemotherapy with precision therapies could improve outcomes, reduce toxicity, and maintain disease control in this TNAC subgroup.

In conclusion, this study provides a comprehensive proteogenomic characterization of TNAC, identifying PI3K/AKT and AR pathway enrichment as key features of invasive progression and revealing metabolic reprogramming as a mechanism of chemotherapy resistance. Our findings suggest that TNAC tumors evade chemotherapy through adaptive survival strategies rather than rapid proliferation, challenging the necessity of standard chemotherapy in this subtype. These insights support a shift toward biomarker-driven therapies primarily targeting PI3K/AKT and AR pathways, with potential consideration for GTPase inhibitors, metabolic interventions, as well as immune therapy, to improve TNAC treatment outcomes.

## Methods

### Tissue Samples

All of the cases diagnosed as invasive breast apocrine carcinoma between 2008 and 2021 in Peking University Hospital were retrieved and reviewed by two senior pathologists. The diagnostic criteria include a minimum of 95% of the tumor displaying apocrine differentiation, tumor cells with a minimum Nuclear:Cytoplasm ratio of 1:2, abundant eosinophilic cytoplasm, distinct nucleoli, and well-defined cell boundaries. Patients with positive expression in ER or PR were excluded. Patients with HER2-amplification were also excluded. A total of thirty-one cases of invasive TNAC were included for proteomic analysis. The FFPE blocks used in this study were obtained from Peking University Cancer Hospital in Beijing, China, with approval from the hospital’s ethical committee under reference number 2020KT113. The cells were stained using H&E, and the expression of protein markers such as ER, PR, and HER2 was evaluated using IHC, as described in a previous publication^9^. Out of the 31 invasive TNAC cases, 10 still contained accompanying DCIS lesions; therefore, invasive, DCIS, and adjacent normal tissues (∼10 μm thick) were carefully dissected from the FFPE blocks on a macroscopic level. In addition, seven patients had received neoadjuvant therapy, and FFPE samples of their needle biopsies taken before chemotherapy were included in the study. The survival of these patients was evaluated using the Miller–Payne grading system.

### Sample preparation for proteomics analysis

Paraffin of FFPE samples was solubilized by xylene, and protein pellets were washed with ethanol. The ethanol was removed completely and the sections were left to air-dry. Samples were lysed with a buffer containing 4% sodium dodecyl sulfate (SDS), 0.1 M Tris-HCl (pH 8.0), 0.1 M DTT (Sigma, 43815), and 1 mM PMSF (Amresco, M145). The collected solution was mixed with pre-cold acetone in a 4:1 ratio. After precipitating the proteins with acetone, they were further washed with cooled acetone. The protein precipitation was then redissolved in 200 μL of 8 M urea solution. The protein concentration was determined by the bicinchoninic acid protein assay (Thermo Scientific, 23227). Samples containing 100 μg of proteins were first reduced with 10 mM dithiothreitol at 56 °C for 30 min and alkylated with 10 mM iodoacetamide at room temperature (RT) in the dark for an additional 30 min. Then, the samples were digested by trypsin using the filter-aided proteome preparation method. Specifically, the samples were transferred into a 30 kD Microcon filter (Millipore, MRCF0R030) and centrifuged at 14,000 × g for 20 min. The precipitate on the filter was washed twice by adding 300 μL of washing buffer (8 M urea in 100 mM Tris, pH 8.0) into the filter and then centrifuged again at 14,000 × g for 20 min. The precipitate was then resuspended in 200 μL of 100 mM NH_4_HCO_3_. Trypsin was added to the filter at a protein-to-enzyme ratio of 50:1 (w/w), and the proteins were digested at 37 °C for 16 h. After this, peptides were collected by centrifugation at 14,000 × g for 20 min and dried in a vacuum concentrator (Thermo Scientific).

### MS analysis

The dried peptide samples were re-dissolved in Solvent A (0.1% formic acid in water) and loaded onto a trap column (100 μm × 2 cm, home-made; particle size, 3 μm; pore size, 120 Å; SunChrom, USA) under a maximum pressure of 280 bar using Solvent A. The samples were then separated on a home-made 150 μm × 12 cm silica microcolumn (particle size, 1.9 μm; pore size, 120 Å; SunChrom, USA) with a gradient of 5-35% mobile phase B (acetonitrile and 0.1% formic acid) at a flow rate of 300 nL/min for 120 min.

The eluted peptides were ionized under 2.2 kV. MS operated under a data-dependent acquisition (DDA) mode. For detection with the Orbitrap Eclipse mass spectrometer, a precursor scan was performed in the Orbitrap by scanning m/z 300-1,500 with a resolution of 60,000. Then, MS/MS scanning was carried out in the Orbitrap by scanning m/z 200-1,400 with a resolution of 15,000. The most intense ions selected under top-speed mode were isolated in Quadrupole with a 1.6 m/z window and fragmented by higher energy collisional dissociation (HCD) with a normalized collision energy of 32%. The maximum injection time was set to 40 ms for full scans and 30 ms for MS/MS scans. Dynamic exclusion time was set as 30 s.

### MS data processing

All MS data were processed in the Maxquant (v 1.6.17.0) platform. The raw files were searched against the human National Center for Biotechnology Information (NCBI) ref-seq protein database (updated on 07-04-2013, 32,015 entries). Mass tolerances were set at 10 ppm for precursor ions and 0.05 Da for product ions. Up to two missed cleavages were allowed. Carbamidomethylation at cysteine was specified as a fixed modification, while acetylation at protein N-terminus and oxidation at methionine were considered as variable modifications. The data were also searched against a target-decoy database to ensure identifications were accepted at a false discovery rate of 1% at both peptide and protein levels. Proteins were classified into protein groups when identified peptides could not differentiate protein isoforms and homologues. The Maxquant-iBAQ labeling-free quantitation method was applied to evaluate protein abundances. Missing values were imputed using minimum values in the corresponding sample. An R package “DEP” (v 1.16.0) was used for data preprocessing and manual examination of data. Before processing, the proteins were filtered to ensure the number of missing values was below half of the smallest sample size condition. The filtered datasets were then log2-transformed and normalized by variance stabilizing transformation to eliminate the effect of different variances among proteins in differential expression protein list.

### GO and KEGG enrichment analysis

Differential protein expressions were analyzed by the R package “limma” (v 3.50.0). Specifically, a separate linear model was fit for each protein, and contrasts were then tested. Empirical Bayes moderation is carried out to obtain a more precise estimate of protein-wise variability. Generally, proteins with an adjusted p-value < 0.05 and a |log2 Fold change| > 1 were identified as differentially expressed proteins. Additionally, R package “clusterProfiler” (v 4.2.2) with default parameters was applied. For the GO enrichment analysis, we focused on enriching “BP” (biological process) and “MF” (molecular function) subontology terms.

### GSEA

For GSEA analysis, R package “clusterProfiler” (v 4.2.2) with default parameters was utilized by the following annotation gene sets from Molecular Signatures Database (v 2023.2.Hs): H (hallmark gene sets), CP (canonical pathways gene sets) and C5 (ontology-gene sets). Differential expression analysis was conducted on gene set expression scores using R package “limma”. Specifically, the KEGG_MEDICUS subset of CP gene sets was evaluated. To reduce the redundancy of highly similar gene sets, we computed the overlap among gene sets using the function “Compute Gene Sets Overlap”. For gene sets with a similarity value larger than 0.8, only one gene set was randomly chosen to be preserved. Pathways with an adjusted P value < 0.01 were identified as significantly altered pathways.

### WGCNA

WGCNA analysis was performed to identify protein modules during TNAC tumor development using R package “WGCNA” (v 1.72-5). 3277 proteins that passed filtering were used for co-expression network construction. Power 10 was selected as the soft threshold as it fits a scale-free topology model well (R2=0.87) while preserving connectivity of 40.72 (Fig. 2d). A signed network was constructed to keep track of the sign of the co-expression information. A topological overlap measure (TOM) was used to evaluate the network interconnectedness, and protein modules composed of highly connected proteins were then identified by hierarchical clustering (Fig. 2d). The modules identified using the blockwise Modules function were further annotated using the compare Cluster function in the R package “cluster Profiler.” This was based on annotation gene sets from Molecular Signatures Database, including H (hallmark gene sets), CP (canonical pathways gene sets), and C5 (ontology-gene sets). The top significantly enriched terms (adjusted p < 0.01) were manually selected to summarize the functions of the modules.

### WES

WES was conducted on six paired invasive and adjacent normal tissue samples from three TNAC patients (ID T001808541, T002002780, T001994386). The exome sequences were efficiently enriched from 0.4 μg genomic DNA using Agilent SureSelect Human All Exon V6 (Agilent USA, Catalog #: 5190-8864)/Agilent SureSelect XT Mouse All Exon library (Agilent, USA, Catalong #: 5190-4643) according to the manufacturer’s protocol. Firstly, qualified genomic DNA was randomly fragmented to an average size of 180-280bp by Covaris LE220R-plus (Covaris, USA). The remaining overhangs were converted into blunt ends via exonuclease polymerase activities. Secondly, DNA fragments were end-repaired and phosphorylated, followed by A-tailing and ligation at the 3’ends with paired-end adapters. DNA fragments with ligated adapter molecules on both ends were selectively enriched in a PCR reaction. After the PCR reaction, libraries hybridize with the liquid phase with a biotin-labeled probe, then use magnetic beads with streptomycin to capture the exons of genes. Captured libraries were enriched in a PCR reaction to add index tags to prepare for sequencing. Products were purified using AMPure XP system (Beckman Coulter, Beverly, USA), libraries were analyzed for size distribution by Agilent 5400 system (AATI) (Agilent, USA) and quantified by real-time PCR (Life Technologies, USA) (1.5 nM). The qualified libraries were pooled and sequenced on Illumina platforms with PE150 strategy in Novogene Bioinformatics Technology Co., Ltd (Beijing, China), according to effective library concentration and the data amount required.

The sequencing results were then aligned to the hg19 reference genome using BWA (v 0.7.8-r455), and the depth was checked by Sambamba (v 0.6.8). Aligned BAM files were processed by SAMtools (v 1.9) to identify the SNP sites and INDEL sites. The SNP sites were annotated using ANNOVAR (v 2017June8) with reference databases, including Refseq, dbSNP, 1000 genome, GO, and KEGG. The final annotations were consolidated into MAF format. Subsequent analyses were conducted based on the samples’ origin using maftools and manual scripts. No threshold for statistical significance was applied.

## Acknowledgment

We thank Weixin Wang and the other team members at Cosmos Wisdom Co., Ltd. in Hangzhou, China, for their contributions to the bioinformatic analysis and graph editing.

## Author contributions

Study conception and supervision: Y.Z., M.L., S.T., T.Hu. Investigation and acquisition of clinical data (sample and patient information collection, pathology review): M.L., T.Hu. Investigation and acquisition of sequencing data: W.S. Data analysis, integration, and interpretation: T.He., F.X., A.Q., Y.L., S.T., W.S., T.Hu., M.L. Supervision of the bioinformatics analysis: Y.Z., S.T., T.Hu Writing: Y.Z., W.S., T.He., F.X. M.L. Reviewing and editing: Y.Z.

## Competing interests

The authors declare that they have no known competing financial interests or personal relationships that could have appeared to influence the work reported in this paper.

## Notes

### Competing Interest Statement

The authors have declared no competing interest.

### Summary of Updates

We performed deeper analyses of proteomic and genomic data in TNAC. Pathway and network results highlight the enrichment of PI3K-AKT-mTOR and AR signaling proteins during tumor progression. Additional analyses of pre- and post-chemotherapy samples reveal changes in metabolic adaptation mechanisms underlying chemoresistance. Figures, text, and therapeutic implications have been updated accordingly.

